# Diversification of the Ruminant Skull Along an Evolutionary Line of Least Resistance

**DOI:** 10.1101/2022.09.13.507810

**Authors:** Daniel Rhoda, Annat Haber, Kenneth D. Angielczyk

**Affiliations:** Committee on Evolutionary Biology, University of Chicago, 1025 E. 57^th^ St., Chicago, IL 60637, USA; Negaunee Integrative Research Center, Field Museum of Natural History, 1400 S. DuSable Lake Shore Dr., Chicago, IL 60605, USA; The Jackson Laboratory, Farmington, Connecticut, USA

**Keywords:** integration, allometry, Ruminantia, Artiodactyla, CREA, macroevolution

## Abstract

Morphological integration is relevant to evolutionary biology and paleontology because the structure of variation within populations determines the ways in which a population can respond to selective pressures. However, understanding the macroevolutionary consequences of morphological integration is elusive because the adaptive landscape is dynamic and population-level constraints themselves evolve. By analyzing a previously published dataset of 2859 ruminant crania with 3D geometric morphometrics and phylogenetic comparative methods, we find that variation within and between ruminant species is biased by a highly conserved mammalian-wide allometric pattern, CREA, where larger species have proportionally longer faces. More tightly integrated species and species more biased towards CREA have diverged farther from their ancestors, and Ruminantia as a clade diversified farther than expected in the direction anticipated by CREA. Our analyses indicate that CREA acts as an evolutionary ‘line of least resistance’ and facilitates morphological diversification due to its alignment with the browser-grazer continuum. These results demonstrate that biological processes constraining variation at the microevolutionary level can produce highly directional phenotypic evolution over macroevolutionary timescales.

## Introduction

Natural selection acts on phenotypic variation in a population. Development structures variation and consequently the ways in which a population can respond to selection (Lande 1979, Lande and Arnold 1983, Alberch 1982, Gerber 2014, Uller et al 2018). The direction with the greatest amount of variation is termed the “line of least resistance” (LLR) (Stebbins 1974, Futuyma et al 1993, Schluter 1996). The LLR represents the direction of greatest potential for evolutionary change because it contains the most variation for selection to act upon and, presumably, because the biological processes underlying its bias easily accommodate evolutionary changes. Populations are expected to evolve in a direct path towards an adaptive peak if selection is aligned with the LLR, but the response to selection will be impeded and diverted towards the LLR if selection is oriented elsewhere (Lande 1979, Arnold 1992, Kirkpatrick and Lofsvold 1992, Björklund 1996). Therefore, the interaction between the adaptive landscape and constraints on variation within a species determine the trajectory of phenotypic evolution.

This interaction operates at the population level. Understanding how constraints within populations scale to explain global patterns of biodiversity is an important avenue for biological research, with the potential to elucidate the microevolutionary mechanisms producing macroevolutionary patterns. Simulations predict that over macroevolutionary timescales, lineages with no constraints on phenotypic variation may explore all areas of morphospace uniformly, but a lineage with a highly constrained phenotype, where traits covary strongly (i.e., high phenotypic integration), will only explore areas of morphospace close to the LLR (Goswami et al 2014, Felice et al 2018). As a result, highly constrained lineages have the potential to evolve more disparate phenotypes than would be expected under a Brownian motion model of evolution, but only along the LLR. Empirically studying the macroevolutionary implications of intrinsic constraints is difficult because the adaptive landscape is dynamic and population-level constraints themselves evolve (Pavlicev et al 2011). Accordingly, studying the relationship between conserved constraints and persistent sources of selection will help us understand the influence of population-level constraints on macroevolution.

In the mammalian skull, a highly conserved pattern of ontogenetic and evolutionary allometry is present, where larger individuals and species have proportionally longer faces (craniofacial evolutionary allometry, CREA; Fig 1) (Radinsky 1985; Cardini & Polly 2013; Cardini 2019). Allometry is historically thought of as a constraint on phenotypic evolution (Gould 1966), and it is insofar as it makes certain trait combinations inaccessible, but allometry also presents an opportunity for novel phenotypes to arise without developmental novelty: extreme phenotypes can arise by ‘piggybacking’ onto relatively labile evolutionary changes in size (Voje et al 2014). In other words, larger, shorter-faced mammal species are improbable, but otherwise out-of-reach long-faced phenotypes are only possible in the largest mammals by exploiting this allometric pattern. CREA may therefore act as an evolutionary LLR and has been hypothesized as such (Marroig & Cheverud 2005, 2010; Cardini & Polly 2013; Cardini 2019, Krone et al 2019).

**Figure 1.**
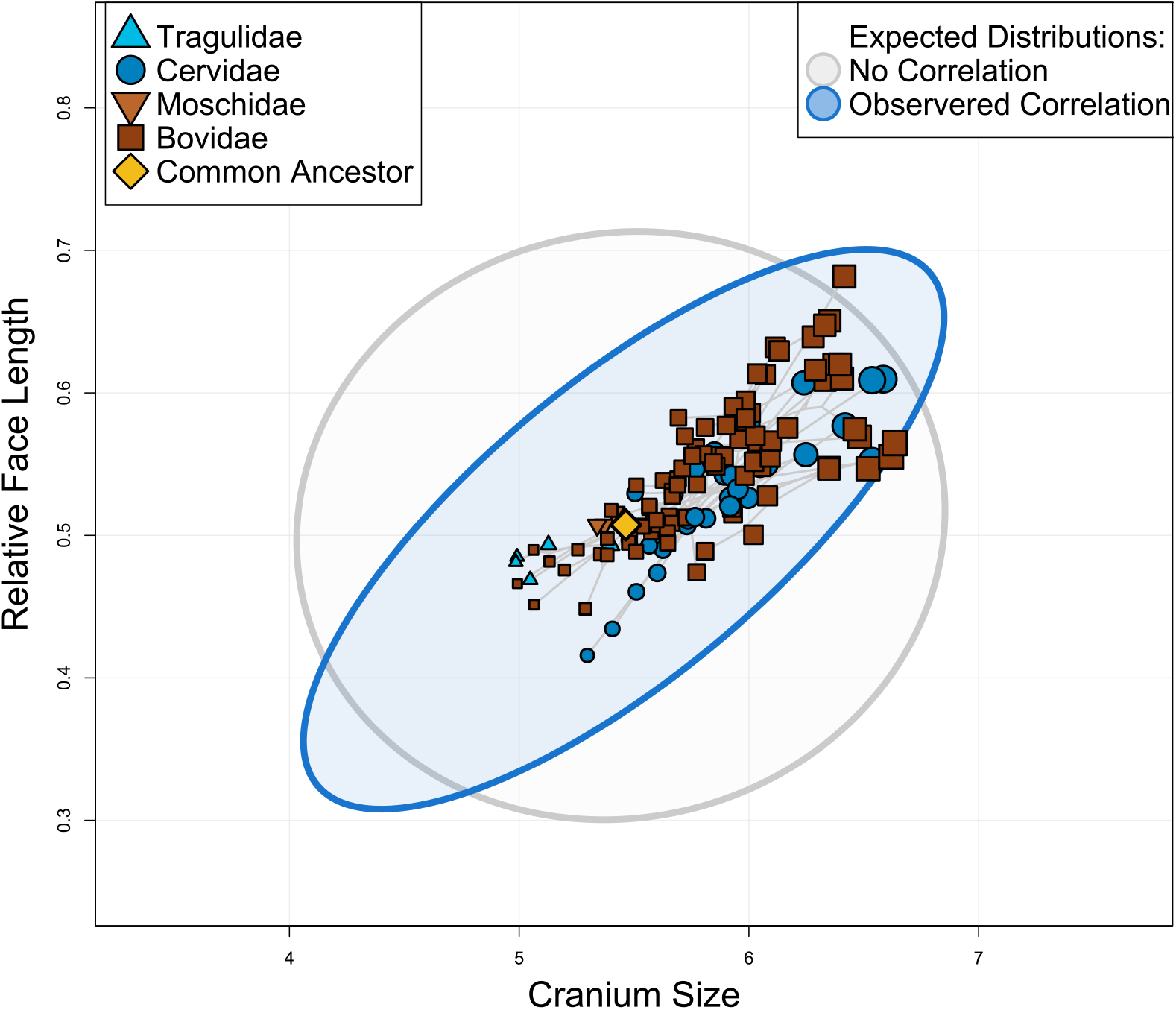
Larger ruminant species have proportionally longer faces, as predicted by CREA (CRaniofacial E-volutionary A-llometry). The gray ellipse represents the 95% confidence interval of the expected distribution of species if there was no allometry of face length (given the observed evolutionary rates), and the blue ellipse represents the expected distribution of species given the observed evolutionary rate matrix. Note that certain ‘extreme’ trait combinations (e.g., very large and long-faced species) are only accessible under correlated evolution, whereas other combinations (e.g., very small, long-faced species) are only accessible under hypothetical uncorrelated evolution.

In ruminant artiodactyls (deer, antelopes, goats, cattle, and relatives), size varies along an ecological axis separating smaller browsing species eating easily digestible foods such as fruit, and larger grazing species subsisting off large amounts of low-growing, high-silica, and nutrient-poor vegetation (Hofmann 1973, Codron et al 2019, with exceptions). Ruminant species span multiple orders of magnitude in body size, from the chevrotains and mouse deer in Tragulidae with masses between 2-3kg, to the gigantic Bovines reaching 1000kg. Haber (2016) found that macroevolutionary patterns of diversification in all ruminants were in part governed by population-level constraints on cranial shape and that within bovids and cervids, species whose structure of variation was better aligned with their clade’s divergence have diverged farther away from their ancestor. A recent study on cranial diversification in Bovidae identified allometry (CREA) as the primary influence on large-scale evolutionary patterns (Bibi & Tyler 2022). In this paper, we present strong evidence that variation at the micro- and macroevolutionary levels is strongly biased by CREA in ruminants, concordant with the definition of an evolutionary LLR, and demonstrate that exploitation of CREA facilitates morphological diversification in directions close to CREA.

## Methods and Materials

To quantitatively test hypotheses about evolutionary constraint, a comparative dataset of intraspecific variation is desirable. We analyzed a previously published dataset of 2859 ruminant crania from 130 species (Haber 2016, with the addition of *Saiga tatarica* and *Megaloceros giganteus*, the extinct Irish Elk) using geometric morphometric methods (Bookstein 1991, Zelditch et al 2012). Landmark definitions are presented in the supplemental materials. All specimens are adults. We split Haber’s (2016) landmark data into two datasets: an interspecific dataset, with a mean landmark configuration representing each species (n = 130, approximately 65% of extant ruminant species), and an intraspecific dataset including only the species represented by at least 27 specimens (n = 49, Haber 2011, 2015). Generalized Procrustes superimpositions were used to remove scale, location, and orientation from the raw landmark data (Rohlf & Slice 1990). Analyses were performed in a phylogenetic context when possible, using a recent molecular phylogeny of mammals (Upham et al 2019) pruned to our taxon sample. This phylogeny did not contain divergence time estimates for subspecies of *Odocoileus*, so we did not divide *O. hemionus* or *O. virginianus* into their subspecies as in Haber (2015, 2016). All analyses were performed in R v4.0.5, primarily using the packages *geomorph, Morpho, mvMorph*, as well as custom scripts available at DR’s Github (https://github.com/danielrhoda) (Baken et al 2021, Schlager 2017, Clavel et al 2015).

### Macroevolution: interspecific metrics & predictions

We estimated the direction of evolutionary allometry (CREA) with a phylogenetic regression of multivariate shape onto log-transformed centroid size in a phylogenetic context using the *procD*.*pgls* function in *geomorph*. We included a species’ subfamily as an interaction term in the model to test if allometric slope significantly differed between subfamilies (Supplemental Fig 1). To visualize patterns of evolutionary variation in the ruminant cranium, we ordinated our interspecific dataset, containing the mean cranium shape for each species, using principal components analysis (PCA, Fig 2). Each individual specimen was projected into this interspecific morphospace. We also ordinated the interspecific dataset both independent of phylogenetic signal (pPCA, with ‘GLS’ and ‘transform’ parameters as TRUE in the gm.prcomp function of *geomorph*) and aligned with phylogenetic signal (PACA, Collyer & Adams 2021), in order to investigate the relative importance of phylogenetic versus ecological signal in the dataset (Supplemental Fig 2-4). An interactive dashboard was created to visualize the distribution of ruminant species in these different ordinations (https://danielrhoda.shinyapps.io/Ruminant_Dashboard/). To aid in the visualization of shape variation, we color-coded the cells of thin-plate spline (TPS) deformation grids according to how much larger or smaller (by area) cells are within the model compared to the reference (undistorted) TPS model. Two-dimensional representations of the 3D morphology from the lateral and dorsal views are presented. Ruminantia is composed mainly of two families, Cervidae (deer) and Bovidae (cattle, antelopes, goats, and relatives), that are characterized by key differences in skull architecture, most notably the presence of annually shed antlers in male cervids but ever-growing horns in both sexes of bovids. In case these clades exploit CREA in different ways, we repeated these ordinations and the phylogenetic regressions of shape on log centroid size using only members of either Cervidae or Bovidae (Fig 3), with the members of the excluded family projected back into the other’s morphospace.

**Figure 2.**
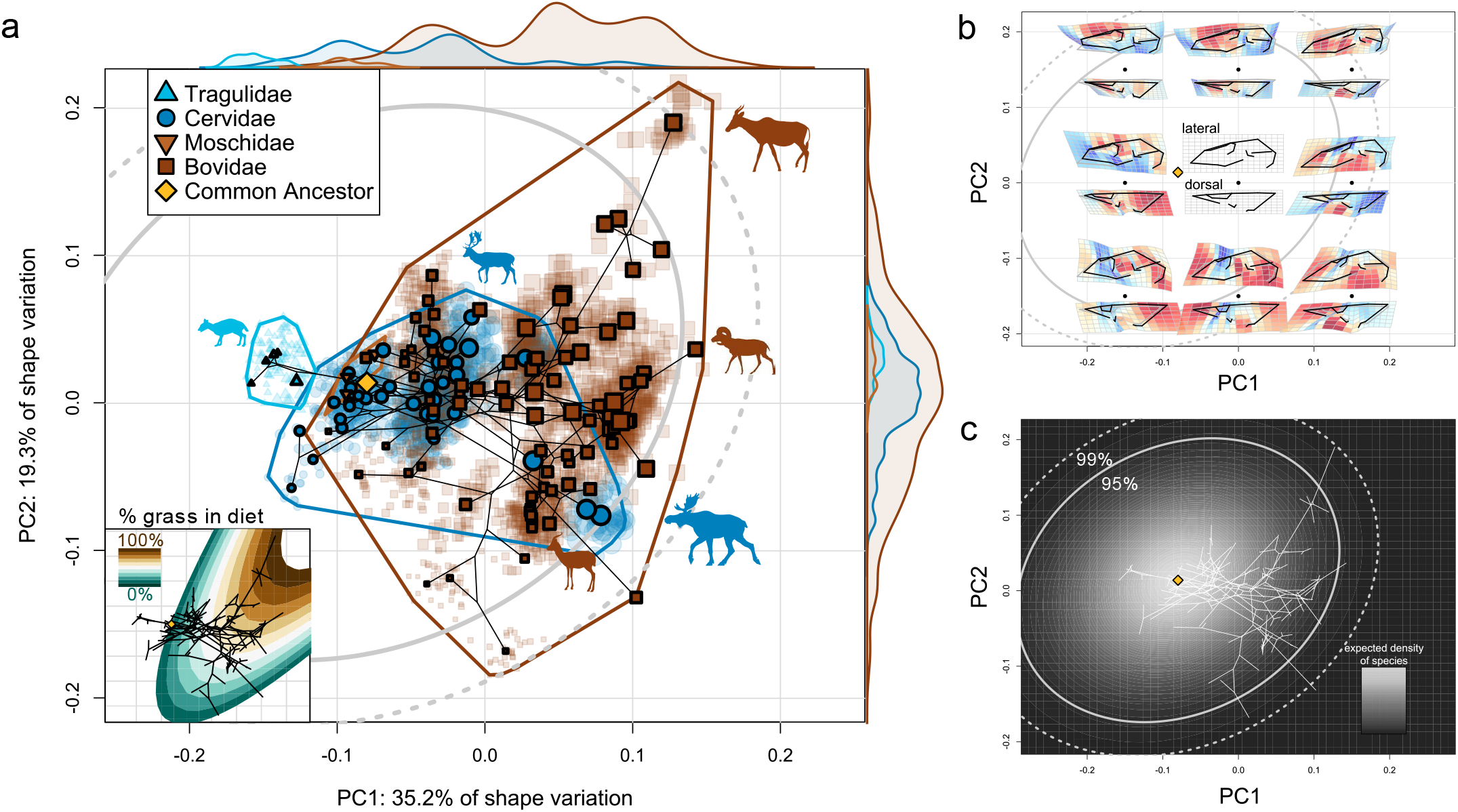
Interspecific PCA morphospace of the ruminant cranium. a) Phylomorphospace of the species’ mean cranial shapes, with individual specimens projected back into the space. Gray ellipses denote 95 and 99% confidence intervals of the probability distribution of species in morphospace given a Brownian motion model of evolution. Size of data point (species) corresponds to centroid size. Probability distributions are shown on the edges of the plot representing the relative densities of each family. (Inset) An ecological surface highlighting variation in % grass in diet, where grazing species consume primarily grass and browsing species consume less grass. Note that variation in % grass in diet is similar to variation in species’ size. b) Shape variation across the morphospace. Each black point represents a shape model at a given coordinate, with its lateral and dorsal views presented above and below, respectively. c) Probability distribution of species in morphospace given our evolutionary rate matrix. Lighter areas denote more-likely areas a species will exist in morphospace given a BM model of evolution. An interactive version of this morphospace (and others) is available (https://danielrhoda.shinyapps.io/Ruminant_Dashboard/).

**Figure 3.**
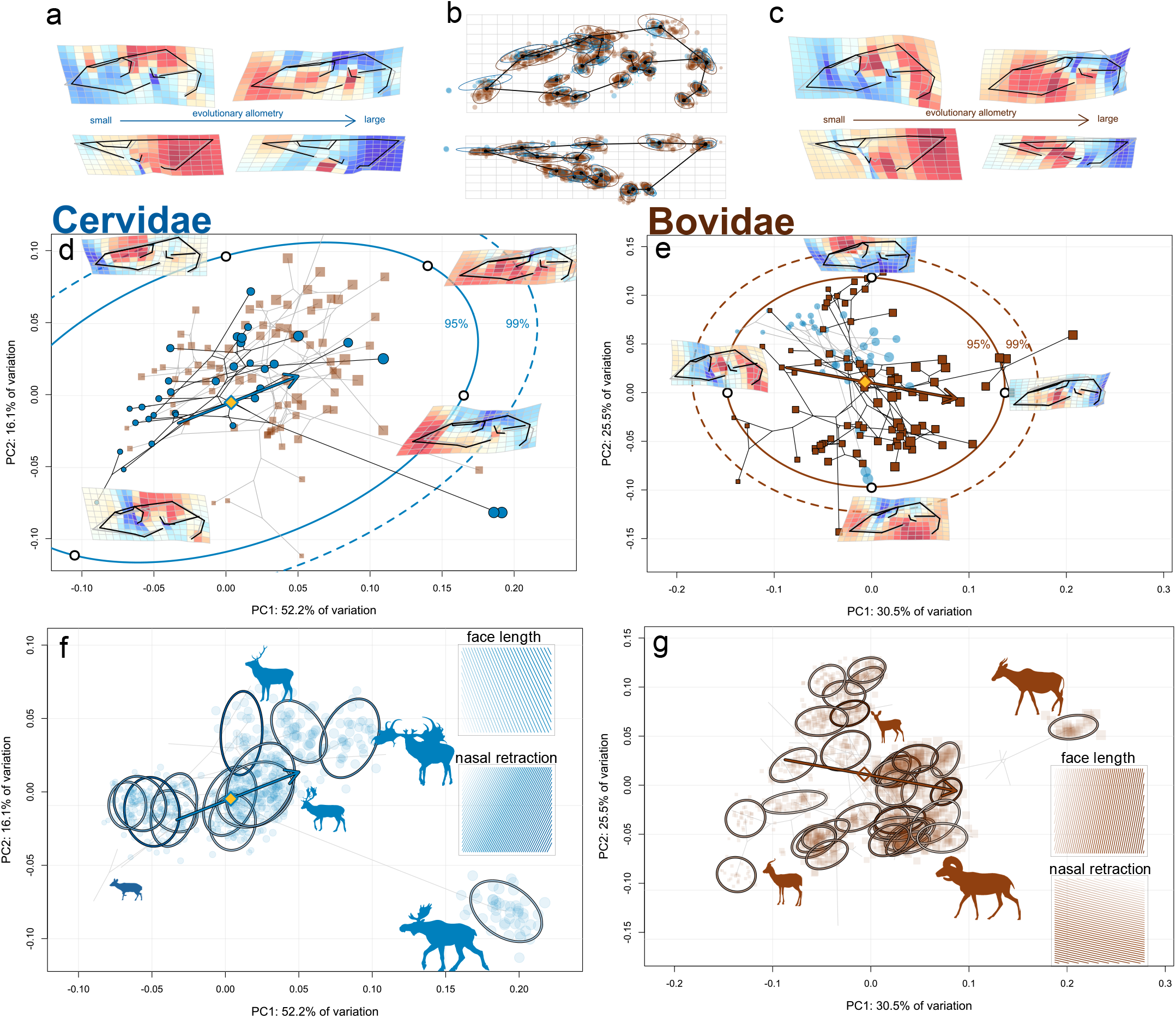
Interspecific morphospaces for Cervidae (left column) and Bovidae (right column). a & Evolutionary allometric trajectories of the cervid (a) and bovid (c) cranium. b) The mean shape of the dataset, with the Procrustes superimposed landmark scatters colored by family and scaled according to species’ centroid size. d & e) Phylomorphospaces of Cervidae and Bovidae, with confidence intervals of the probability distribution of species in morphospace, as in Figure 1. Arrows denote allometric trajectories, as in panels f and g. f & g) Family-specific morphospaces with individual specimens projected into them and ellipses for each species in the intraspecific dataset. The strength of ellipse color corresponds to the species’ V_proj_ value. (inset, top) Face spaces: variation in relative face length across the morphospace, where darker areas have longer faces. (inset, bottom) Variation in nasal retraction, where darker areas have farther nasal retraction.

To examine the relationship between cranial morphology and the browser-grazer continuum in ruminants, we collected percent-grass-in-diet (a proxy for a species’ location on the browser-grazer continuum; Janis 1995) from Codron et al. (2019) for a subset of our interspecific dataset (n=98). We fit a second-degree polynomial surface to the empirical percent grass data and PC1 and PC2 scores (cumulatively describing 54.5% of total variation) using the *surf*.*ls* and *trmat* functions in the *spatial* R package (Venables & Ripley 2002). Exclusively sampling naturally occurring morphologies, rather than a grid of theoretical morphologies, has been shown to produce accurate surfaces (Smith et al 2021). This ecological surface aids in the visualization of variation in diet as it relates to variation in morphology (Fig 2). For the family-specific morphospaces, we produced surfaces corresponding to variation in relative face length and nasal retraction (discussed further below). These surfaces were constructed from a grid of theoretical morphologies because face length and nasal retraction indices can be readily computed from theoretical shapes across the morphospace. Face length was calculated as the distance between the anterior tip of the premaxilla (landmark 8) and anterior orbit (landmark 18) divided by total length of the skull following Bibi & Tyler (2022). Nasal retraction was calculated as the distance between the anterior tip of the premaxilla (landmark 8) and the anterior tip of the nasal (landmark 22) divided by skull length. All surface fits were significant (i.e., the empirical PC1/2 scores significantly covaried with the trait data; p-values < 0.001).

In an integrated phenotype, accessible areas of morphospace are limited, but evolutionary rate is unaffected (Goswami et al 2014, Felice et al 2018). Goswami, Felice, and colleagues (2014, 2018) ask us to consider a “fly in a tube” model of the evolutionary consequences of integration, where a fly (phenotype) can ‘zip’ around a tube (morphospace) at any speed (evolutionary rate), but only within a tube of a specific shape, dictated by integration pattern. Under Brownian motion, evolutionary rates are held constant, and variance is proportional to time. Therefore, given an observed rate of phenotypic evolution, we can predict whether or not integration is facilitating or obstructing phenotypic evolution by testing whether disparity is proportional to rate. In some recent studies this was tested by regressing per-landmark Procrustes variance onto evolutionary rate and observing the slope of the relationship (e.g., Felice & Goswami 2018, Bardua et al 2019, Fabre et al 2020, although see Cardini & Marco 2022). In this study, our analytical units are species, not landmarks, so we took a different approach to test the ‘fly in a tube’ model. First, we fit PC1-5 scores of the interspecific morphospace to a Brownian motion model of evolution to compute an evolutionary rate matrix of PC scores (function *mvBM* in *mvMorph*; Clavel et al 2015). We could not compute a rate matrix for all PC axes because of computational constraints, so we chose to only include the ‘meaningful’ PC axes *sensu* Bookstein (2014). This rate matrix, **C**, contains the evolutionary rates of each PC along the diagonal, and pairwise co-evolutionary rates elsewhere. Assuming Brownian motion, **C** defines a multivariate normal probability distribution of species distribution in morphospace, centered at the estimated ancestral shape. In other words, if we were to simulate random walks of PC scores a large number of times from the ancestral shape given our observed evolutionary rate matrix, the highest density of species would be concentrated around the ancestral shape with lower densities farther away from the ancestral shape (Fig 2c). We calculated 95% and 99% confidence intervals for this probability distribution and overlayed it on our morphospaces to visualize areas in morphospace where lineages defy expectations. If CREA is being exploited as an LLR and facilitating diversification, we would expect some lineages to evolve beyond the confidence intervals (i.e., “fly” to the ends of the “tube”), but only by following the trajectory defined by CREA.

### Microevolution: intraspecific metrics & predictions

The question we address in this paper is whether CREA is being exploited as an evolutionary LLR in ruminant artiodactyls, and how this influences morphological diversification. Schluter (1996) tested for the presence of an LLR by successfully predicting that the direction in which species diverge from their ancestors should be biased towards the principal direction of intraspecific variation (PC1 of an additive genetic covariance matrix, **G**, or a phenotypic covariance matrix, **P**, used as substitute for **G**; Cheverud 1988) and that species with greater alignment between the direction of their variation and divergence should diverge farther. Statistically testing the congruence between evolutionary divergence and PC1 of **G** or **P** matrices has remained the standard means for documenting LLRs since Schluter’s seminal work (e.g., Boell 2013, Polly & Mock 2018, McGlothlin et al 2018, Marroig & Cheverud 2005, Renaud et al 2006, Fasanelli et al 2021). Considering PC1 as the *de facto* LLR is intuitive because, in the presence of a strong LLR, responses to selection should be channeled into the direction of the LLR, consequently forming the major axis of interspecific phenotypic variation. However, PC axes are the major axes of variation irrespective of any specific generative process, and their sensitivity to sample details means that there is no guarantee that a given axis will capture variation uniquely associated with a process of interest (Houle et al 2002; Mezey & Houle 2003). A better approach, we argue, is to directly measure the axis of variation associated with a hypothesized LLR (i.e., some developmental process, in our case CREA), and then interrogate its relationship with population-level and evolutionary variation. So as not to place undue importance on an arbitrary axis of variation, we adopted the ‘projected variance’ approach used by Hunt (2007) in this paper. The **P** matrix for each species in the intraspecific dataset was projected onto the evolutionary allometric axis, described as a unit-length vector of allometric coefficients, using the following equation:

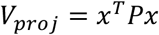

where **P** is the species’ phenotypic variance-covariance matrix (of superimposed landmark data) and **x** is the vector of allometric coefficients (T indicates transpose). Projecting **P** onto **x** gives us a number, V_proj_, describing the proportion of shape variation in **P** described by the projected vector. A higher V_proj_ value indicates that the species’ variation is more biased in the direction of CREA. This measure is equivalent to the ‘evolvability’ metric *sensu* Hansen & Houle (2008) of a **P** matrix given a certain direction. To examine how aligned population-level variation is with CREA versus other major axes of evolutionary variation (Fig 4a), we compared the distribution of V_proj_ values from CREA to 1) a distribution of V_proj_ calculated from random vectors (for each species, the average of 499 random vectors); 2) V_proj_ of the direction of divergence from the species’ subfamilies’ inferred ancestor; and 3) the first two eigenvectors from alternative ordinations of the between-species covariance matrix (PCA, pPCA, PaCA). V_proj_ was also computed for bovids and cervids separately using their clade-specific ordinations and evolutionary allometric trajectories (Fig 4b & c).

**Figure 4.**
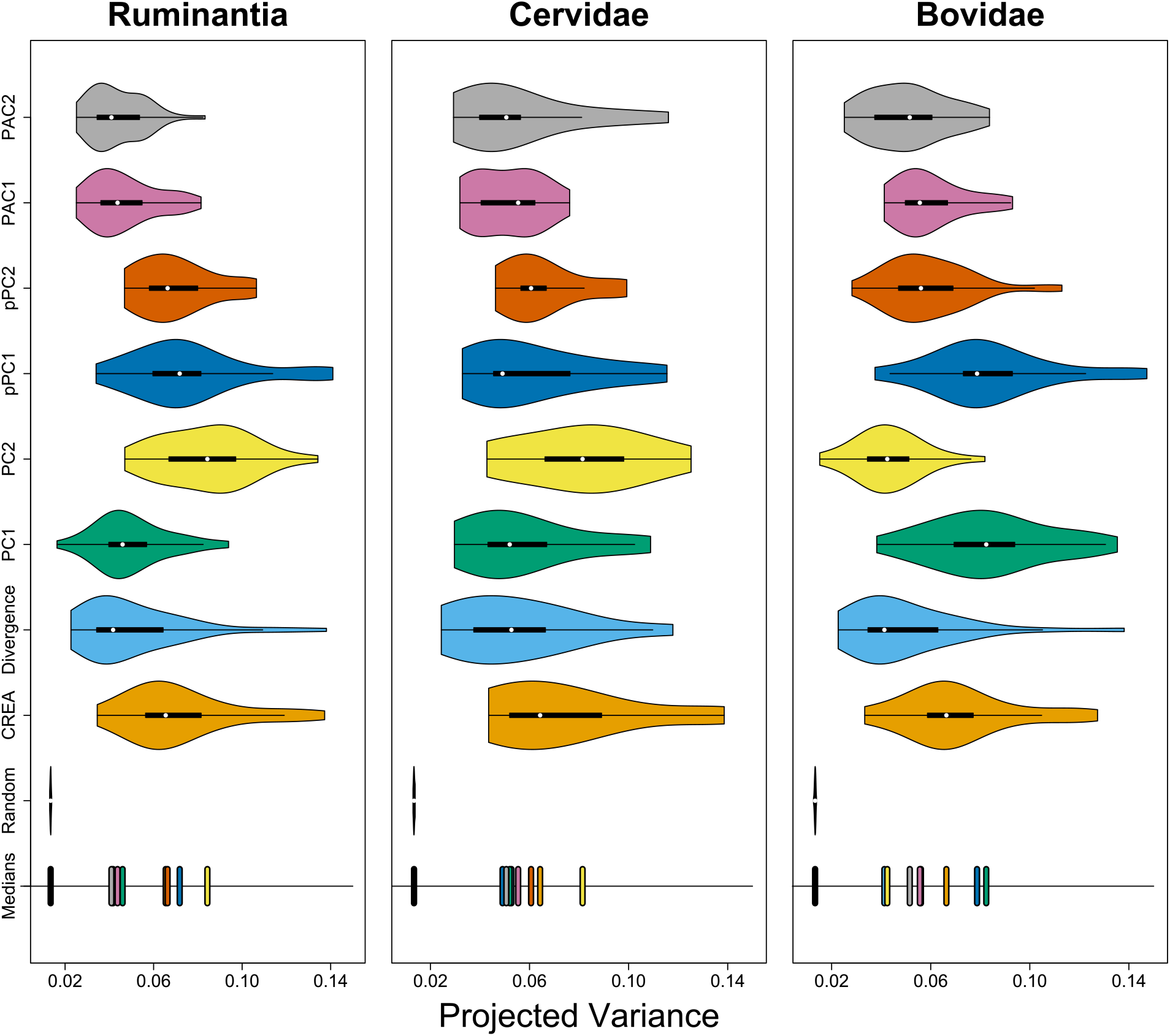
Distributions of species’ projected variance values for different directions of evolutionary variation for all of Ruminantia (left), Bovidae (middle), and Cervidae (right). Axes of evolutionary variation for Cervidae and Bovidae are calculated using only members of the respective clade. For example, PC1 & PC2 in the cervid and bovid panels are the axes presented in Figure 3. The positions of medians of the distributions are at the bottom of each plot.

If CREA is being exploited as an LLR, species with **P** matrices closely aligned with CREA (higher V_proj_) should have diverged farther from their ancestor than less-aligned species, and species that have diverged in the direction of CREA should have diverged farther from their ancestor than species diversifying in other directions. It is also possible that species with strongly biased **P** matrices may already lie on an adaptive peak and are under stabilizing selection, so as long as species with marginal bias towards CREA have not diverged farther from their inferred ancestor than species with strong bias, CREA may have been exploited as an LLR. We tested whether the direction of species’ divergence was biased by CREA by comparing the magnitude of a species’ divergence from its inferred ancestor to 1) the species V_proj_ value of CREA, the species V_proj_ of the direction of its divergence (the amount of variation a population has in the direction it evolved from), and magnitude of morphological integration (Fig 5); and 2) the angle between the direction of divergence and direction of CREA (Fig 6). Morphological integration was quantified as the standardized effect size of relative eigenvalue variance of each species’ **P** (Z_rel_, *integration*.*Vrel* function in *geomorph*; Pavlicev et al 2009, Haber 2011), which has been shown to perform better than other integration metrics under controlled simulations (Watanabe 2022, Conaway & Adams 2022). Morphological divergence was calculated as the Euclidean distance between the PC scores of a species and the PC scores of its subfamily’s inferred ancestor (as in Haber 2016). The most recent common ancestor of the subfamily was used rather than the position of the immediate ancestral node because our dataset does not contain all ruminant species and the immediate ancestral node’s position is reliant on only the positions of its two descendants. The angle between divergence and CREA was computed by calculating the difference in coordinate values between a species’ landmark configuration and that of its subfamily’s inferred ancestor, and then taking the angle between this vector and the vector of evolutionary allometric coefficients. Phylogenetic generalized least-squares regressions (Grafen 1989, Martins & Hansen 1997) were used to measure the associations between morphological divergence and the angle of divergence relative to CREA, and between morphological divergence and integration and V_proj_.

**Figure 5.**
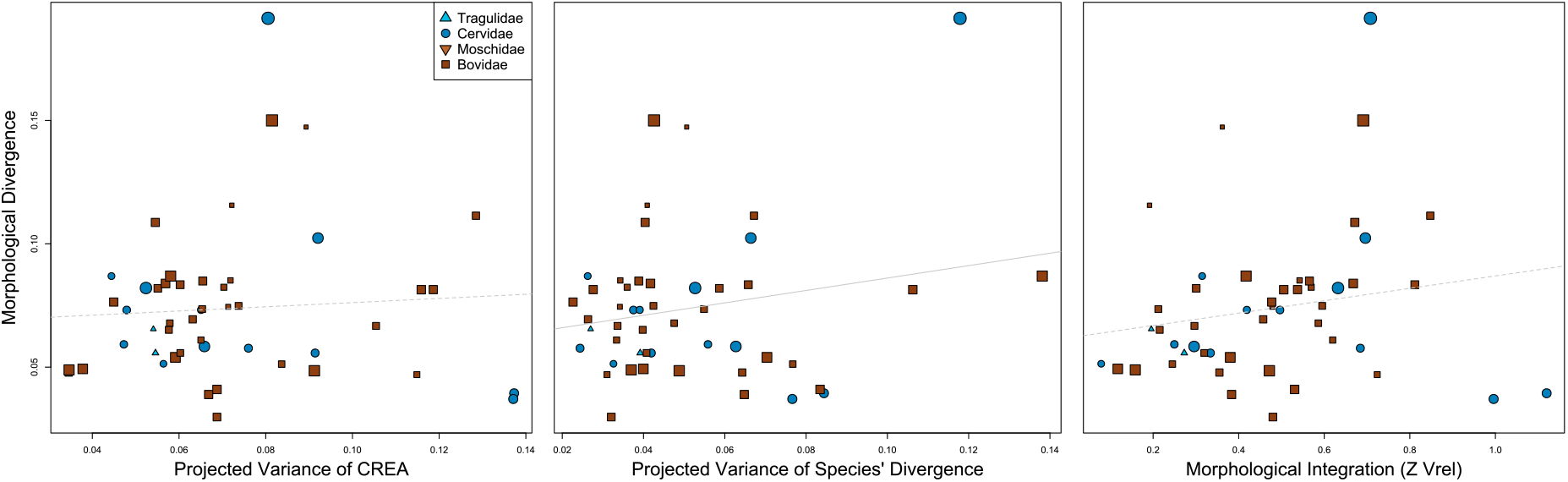
Scatterplots of properties of **P** and the magnitude of divergence from an ancestor. Each point represents a species, with the point size corresponding to centroid size.

**Figure 6.**
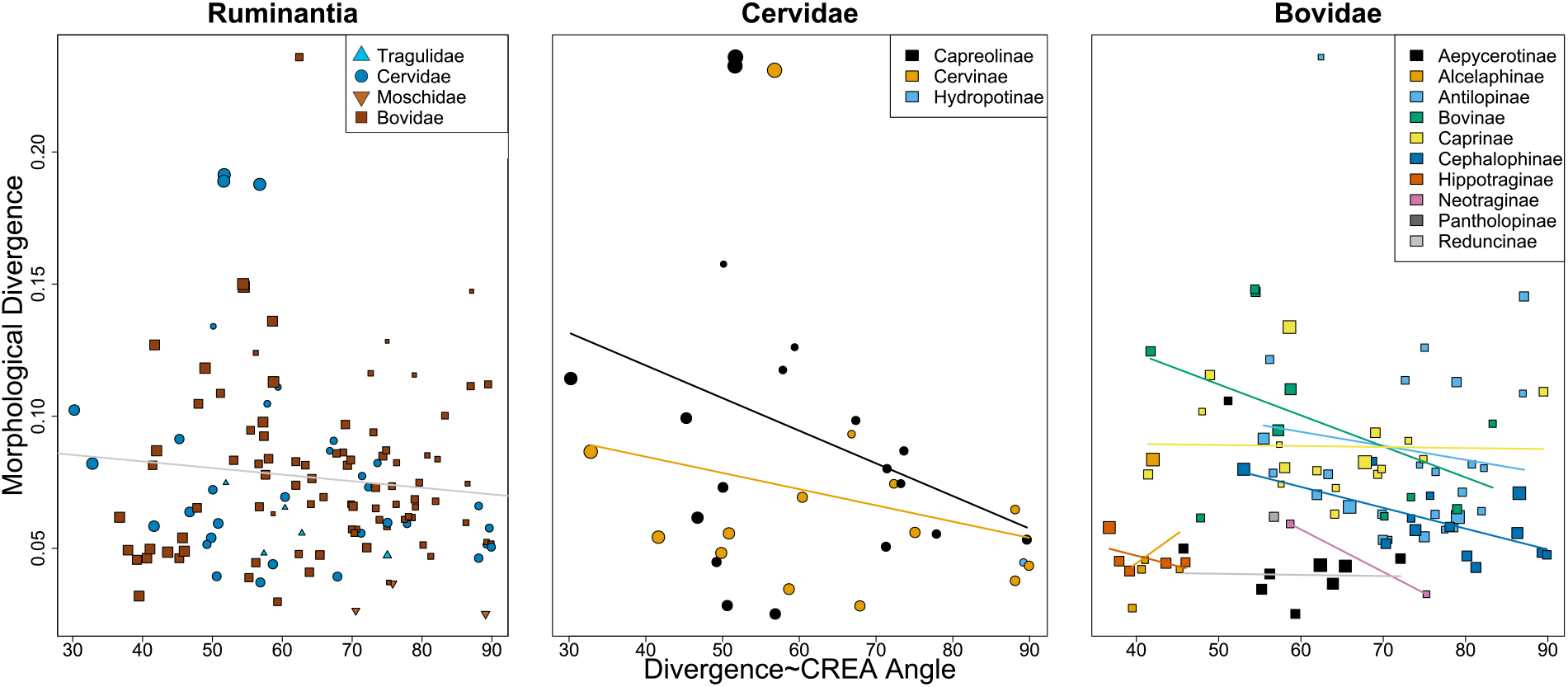
Scatterplots of the magnitude of morphological divergence and the angle between the direction of divergence and CREA. All ruminants in our dataset (left), Cervidae (middle), and Bovidae (right). Each point is a species and point size corresponds to centroid size.

## Results & Discussion

### CREA Governs Macroevolutionary Trends in the Ruminant Skull

A strong relationship between size and skull shape between species was found, where larger species have longer faces (Fig 1, p-value < 0.001, R^2^ = 0.196, Z = 5.927, F = 31.191). The slope of evolutionary allometry varied between subfamilies significantly (p = 0.012), but in each subfamily larger species had longer faces (Supplemental Fig 1).

The major axes of interspecific variation preserve a clear trend in size (Fig 2), where larger species are located at the positive ends of PC1 and PC2 (describing 35.2% and 19.2% of shape variation, respectively). The first PC captures variation in the relative length of the face and lateral expansion of the orbits (related to the characteristic tubular shape of bovid orbits). The second PC describes relative face length variation as well, but also captures variation in the posterior retraction of the nasals relative to the premaxilla and overall slenderness of the skull (shapes closer to the positive end are more gracile with less nasal retraction and longer faces). Distributions of the density of specimens (per family, accounting for different sample sizes of species) show a clear phylogenetic structure to PC1. The shapes of the species distributions are similar along PC2, but more diverse families extend farther in morphospace. Our estimated common ancestor is short-faced and small, observed at a negative PC1 score and near-zero PC2 score. Consequently, many of the ruminant species that extend beyond our 95% confidence intervals are large bovids with positive PC1 scores. Most notably, the hartebeest, *Alcelaphus buselaphus*, explores extreme PC1+ and PC2+ values and has the longest face of the species in our dataset. Most small ruminant species lie near the common ancestor’s shape, but the dik-diks (genus *Madoqua)* have more negative PC2 scores than any other ruminant. The cranial shapes of dik-diks are unlike the other small ruminants in having greater nasal retraction. Regardless, all small ruminants, the tragulids, moschids, and browsing cervids and bovids have relatively shorter faces than their massive, long-faced, relatives. Moose, *Alces americanus* and *A. alces*, are the only cervids to extend beyond the 95% confidence interval of the interspecific morphospace and deviate far from all other cervids other than *Megaloceros giganteus*, the Irish elk. Unsurprisingly, *Alces* and Irish elk are by far the two largest cervids in our dataset. Our pPCA and PACA morphospaces both show clear size trends dominating the major axes of variation, further strengthening our findings (Supplemental Fig 2-4). Importantly, lineages that explore the most distant regions of morphospace are particularly short-faced and small or long-faced and large.

We recreated the interspecific morphospace for Cervidae and Bovidae individually to understand potential differences between the clades in their evolutionary patterns and exploitation of CREA (Fig 3). Size explained large amounts of variation in both Cervidae (p-value < 0.001, R^2^ = 0.354, Z = 4.243, F = 17.506) and Bovidae (p-value < 0.001, R^2^ = 0.167, Z = 5.898, F = 17.287) (Fig 3a,c). A majority (52.2%) of cranial variation in the cervid skull is explained by the first PC, which describes the length of the face and nasal retraction (both increase towards positive end, Fig 3d,f). Other than *Alces*, interspecific differences in cervid skull shape are highly eccentric in the direction of CREA but cervids occupy a restricted area of morphospace compared to Bovidae. The first PC of bovid morphospace describes variation in relative face length and height of the cranium, and is well-aligned with the family’s evolutionary allometric trajectory (Fig 3e,g). The second PC describes nasal retraction, with *Saiga* at the negative end and the frugivorous duikers (Cephalophini) at the positive end. The morphospace is strikingly similar to the morphospace of Bovidae presented by Bibi and Tyler (2022), who argued that allometry was the primary influence on bovid skull diversification. We corroborate this finding. There is a clear size trend to the major axis of evolutionary variation in the bovid cranium, and it is this axis where the large, grazing tribe Alcelaphini and the small, browsing tribe Neotragini diversify farther than expected given their observed evolutionary rate. Within both families, patterns of intraspecific variation are biased in the directions of their allometric axis (Fig 3f,g), with exceptions (e.g., *Alces*, but note that the direction of *Alces’* ellipse is congruent with the branch separating it from the rest of deer).

*Alces* is the farthest lineage along PC1+ in cervid morphospace by far, extending well beyond the 99% confidence interval. The Saiga antelope, a bovid species with an unusually high degree of nasal retraction, is the only other taxon near *Alces*. Both *Saiga* and *Alces* have probosces overhanging their mouths, but the adaptive significance of these structures seems to differ between the species (Clifford and Witmer 2004a, b). The Saiga lives in the deserts of the Central Asian steppe and uses its enlarged proboscis to regulate the entry of dust during respiration (Clifford and Witmer 2004a), but Boreal moose, on the other hand, often forage semi-aquatically for sodium-rich vegetation that is required for antler growth and is absent in its terrestrial environment (Aho and Jordan 1979). It is hypothesized that moose use their proboscises as a valve to prevent the flow of water into the airway (Clifford and Witmer 2004a). Mating behaviors, either loud vocalizations amplified by the proboscis (*Saiga*, Frey et al 2007) or ‘lip-curling’ in *Alces* (Marquez et al 2019) serve as additional explanations for nasal retraction. The rest of cervid species and their constituent specimens are restricted to a scatter biased in the positive direction, highly congruent with the direction of CREA. It would be reasonable to assert that moose do not adhere to CREA because the branch leading to *Alces* is nearly orthogonal to CREA in this projection of tangent space, but this is partly misleading – *Alces* has the longest face and is the largest extant cervid genus, and the primary difference between it and other cervids is the unusual degree of nasal retraction. Notably, *Cervalces* (not present in our dataset), an extinct relative of *Alces*, does not share *Alces*’ peculiar nasal retraction (Breda 2008), nor does the Irish elk, which is comparable in size to *Alces*. The modest size of *Saiga*, the presence of more nasally restricted morphologies in *Madoqua*, and the presumptive recent evolution of nasal restriction in *Alces* suggested by *Cervalces* all suggest that this morphotype is not necessarily related to size.

### The Adaptive Significance of CREA

It is unclear whether the exceptionally long-faced crania of large grazing species like the Alcelaphini are 1) a direct result of selection for crania better adapted for grazing, which coincidentally happen to be long-faced and easily evolvable under the clade-wide allometric constraints, or 2) are agnostic to selection for foraging ecology, with longer faces passively evolving during increases in body size associated with grazing (or decreases in size associated with browsing). The latter seems unlikely considering that the face is intimately involved in food ingestion and processing. For example, a proportionally longer face would allow larger amounts of plant material to be processed at once and more room for cranial musculature to aid in mastication (Endo et al 2002, Clauss et al 2008a), which may be advantageous for large grazing species that eat gritty, fibrous materials. A longer face also keeps the eyes farther away from the ground, both protecting the eyes and providing better detection of predators (Janis 1995, Clauss et al 2008b). Regardless, as long as longer faces are not maladaptive for grazing ecologies and shorter faces maladaptive for browsing, CREA would not obstruct evolution along the browser-grazer continuum and instead facilitate diversification of skull form. In either of our two probable scenarios, foraging ecology provides adaptive value for evolutionary changes in size, and thus motivation for ruminant cranial morphology to fly to the ends of its tube.

Although we have focused on the significance of a few exceptional lineages that defy the predictions of Brownian motion, a majority of ruminant species have intermediate face lengths and sizes, and cluster at neutral PC2 values. Most ruminants are not obligate browsers or grazers but are facultative ‘mixed feeders’ who opportunistically forage on a combination of different plant materials. Multiple studies have suggested that lineages change their diets towards mixed feeding when faced with environmental turbulence (Badgley et al 2008, Codron et al 2008, DeMiguel et al 2008, DeMiguel et al 2010). Central locations in interspecific morphospace (Fig 2) may therefore be consistent with the concept of a “net adaptive peak” (Polly 2020). In a biological system with functional tradeoffs, the optimal phenotype (adaptive peak) is not necessarily one that maximizes a single function, but a neutral phenotype able to adequately perform multiple functions, with the relative importance of competing functions determined by the environment. The high density of cranial forms lying in central locations of morphospace, suitable for browsing *and* grazing, solidifies this claim. In Ruminantia, mixed feeding originated around the Oligocene-Miocene transition and triggered a period of taxonomic and ecological diversification (Cantalapiedra et al 2014). Grazing behavior emerged near the Mid-Miocene climatic optimum and triggered another radiation during the expansion of C_4_ grasses, but transitions between browsing and grazing (exclusively through mixed feeding) remained common (Cantalapiedra et al 2014). The data presented in this paper suggest the evolutionary lability of ruminant diet through the Neogene, and therefore their evolutionary success, may in part be explained by the alignment of CREA and this ecological axis. As the environment changes, ruminant lineages can simply slide along an allometric axis to adapt to different relative amounts of browsing and grazing. Contrasting these observations in Ruminantia with perissodactyls may help explain perissodactyls’ declining diversity throughout the Cenozoic: is the covariance structure of the perissodactyl skull less aligned with the browser-grazer continuum, obstructing ecological transitions and thus limiting diversification?

### Population-level Variation is Biased Towards CREA

As discussed above, macroevolutionary patterns reveal a role for CREA in influencing morphological diversification, but how is this reflected at the population level? For CREA to qualify as an evolutionary LLR, intraspecific variation should be biased in the direction of CREA, species should diverge from their ancestor primarily in the direction of CREA, and species that align more closely with this direction should diverge farther. We find that intraspecific variation is consistently biased in the direction of CREA. CREA explained 3.5-13.7% of variation within species, which was much greater than the variation explained by the distribution of random vectors (Fig 4), suggesting that allometry is indeed a bias on intraspecific variation. The major axes of evolutionary variation explained similar amounts of variation as CREA; only PC2 of the interspecific morphospace explained considerably more variation, indicating that intraspecific variation is as aligned with CREA as it is with the other major directions of evolutionary variation (Fig 4). Population-level variation was even more biased towards CREA than the direction in which the species diverged.

Phylogenetic GLS regressions found no significant relationship between V_proj_ of CREA and the magnitude of species’ divergence from their subfamily’s ancestor (Fig 5a; p-value = 0.952, F=0.003), but there was a significant relationship between the magnitude of divergence and amount of variation in **P** explained by divergence (Fig 5b; p-value = 0.034, F = 4.76). As mentioned above, species with their structure of variation aligned with CREA may already lie on adaptive peaks and have no incentive (or ecological opportunity) to diverge far. The lack of ruminant species with low V_proj_ values that have diverged far is consistent with our predictions. The magnitude of divergence and morphological integration within a species had a slight positive relationship (Fig 5c; p-value = 0.123, F = 2.47). *Odocoileus virginianus* and *O. hemionus* were the most tightly integrated species and most biased towards CREA but they did not diverge far from their ancestor. Their high Z_rel_ and V_proj_ values may be due to the presence of subspecies if the axis of variation separating subspecies is similar to the primary axes of variation within the subspecies. When removing *Odocoileus* we find a strong positive relationship between integration and divergence (p-value = 0.011, F = 7.04), and a slightly stronger but insignificant relationship between bias towards CREA and divergence (p-value = 0.315, F = 1.03). The positive relationship between integration and divergence suggests that highly eccentric variation is contributing to morphological diversification, but that this eccentric variation is not always due to allometry. Species that diverged in the direction of CREA diverged farther than species diverging elsewhere (p-value = 0.012, F = 6.44) (Fig 6). This relationship is present in cervids but not significant (p-value = 0.087, F = 3.12) (Fig 6b) and less so in bovids (p-value = 0.181, F = 1.82) (Fig 6c). There seem to be certain ruminant subfamilies that preserve this relationship, such as Bovinae, Antilopinae, Cephalophinae (Fig 6c), and the cervid subfamilies Capreolinae and Cervinae (Fig 6b), but out of the six ruminant subfamilies with at least 10 species (the previously mentioned subfamilies as well as Caprinae), the relationship is only significant in Cephalophinae (p-value = 0.012, F = 9.18). Note that in hyperdimensional spaces like the one here (k=75), a distribution of random angles will be normally distributed around 90 degrees with a lower standard deviation than lower-dimensional spaces. The angles of divergence and CREA was significantly different than a distribution of 10000 random k-dimensional angles (two-sided Kolmogorov-Smirnov test, p-value < 0.001) (Supplemental Fig 6), meaning that morphological divergence was more aligned with CREA than would be expected by chance.

### Micro- and Macroevolutionary Consequences of CREA as a Line of Least Resistance

Taken together, the relationships between the structure of variation (**P**) and the direction and magnitude of divergence reinforce the argument that CREA is being exploited as an evolutionary line of least resistance. At the macroevolutionary level, cranial morphology of ruminants diversified farther than expected under our observed evolutionary rates assuming Brownian motion, but only in the direction of CREA. At both micro- and macroevolutionary scales, highly integrated cranial variation led to the exploration of greater ranges of morphospace.

Craniofacial elongation is characteristic of postnatal growth within mammalian species. Our analyses suggest that this well-known allometric mechanism, where the growth of the face outpaces the braincase, defines an evolutionary LLR and governs both population- and clade-level patterns of morphological diversity in Ruminantia. Haber (2016) demonstrated that in the ruminant skull evolutionary divergence is well-aligned with **P** for most species, and that the better the alignment the farther species diverge from their clade’s ancestor. Our reanalysis of Haber’s landmark dataset expands these findings and implicates a mammalian-wide allometric pattern as the biological process likely underlying these consequential intrinsic constraints. There is still, however, substantial flexibility within the constraint imposed by CREA. The structure of **P** has evolved throughout the history of Ruminantia (Haber 2015), possibly due to other factors influencing skull form such as ornamentation. Despite these reorganizations in covariance structure, CREA consistently explains as much or more intraspecific variation as the other major directions of evolutionary divergence (Fig 4). It is unrealistic to expect that covariance structure is perfectly conserved throughout a clade’s history (Arnold et al 2008), especially considering that variation in directions other than CREA needs to be maintained to respond to other selective pressures, but the analyses presented herein demonstrate that certain biological processes at the population-level can impose constraints that dominate macroevolutionary trends even though covariance structure evolves.

The relationship between phenotypic integration and disparity is a highly active area of research with little agreement in empirical results. Studies have found positive (e.g., Navalón et al 2020, Hedrick et al 2020), negative (e.g., Goswami & Polly 2010, Felice & Goswami 2018), and insignificant (Watanabe et al 2019, Marshall et al 2019, Bardua et al 2019, Bon et al 2020, Bardua et al 2020, Martín-Serra & Benson 2020, Rhoda et al 2021) relationships between different metrics of the magnitude of morphological integration and disparity at different biological levels. These conflicting results may be because the directions of selection and integration are usually semi-independent (although see Pavlicev et al 2011), but their interaction determines a population or clade’s ability to diversify in certain directions (Lande & Arnold 1983). In reality there are likely numerous selective pressures in different directions, and any given structure of integration is impeding responses to some of these pressures and facilitating responses to others. A universal relationship between the strength of integration and disparity is therefore unlikely, and the relationship should depend on the relative strengths of the different selective pressures and their alignment with **P**s within a clade of interest. In the case of Ruminantia, size dominates cranial variation through allometric scaling of facial length and is related to foraging ecology (Bodmer 1990, Pérez-Barbería & Gordon 2001), signaling congruence between the direction of selection and integration. Accordingly, here we generally find positive relationships between morphological integration and disparity at both micro- and macroevolutionary levels. If not for the fortuitous alignment between a highly conserved allometric pattern and selection, this pattern is unlikely.

### CREA and Mammalian Evolution

Craniofacial elongation characterizes allometric patterns within and between mammalian species of nearly all placentals (Cardini & Polly 2013, Tamagnini et al 2017, Cardini 2019, Marcy et al 2020), probably marsupials (Cardini et al 2015, Newton et al 2021), and some groups of non-mammalian synapsids (pelycosaurs, gorgonopsians, and possibly others, Krone et al 2019). The CREA pattern is highly conserved and deeply rooted in mammalian evolution. Does CREA act as an evolutionary line of least resistance in other clades? And if so, how ubiquitous is exploitation of CREA in mammals? We speculate that exploitation of CREA is probably common, or at least not rare, considering the myriad ways modifying relative length of the face can confer adaptive value. For example, a relatively shorter face generates stronger bite forces because of lever mechanics, which has consequences for feeding ecology. Phyllostomid bats contain the highest dietary diversity of any mammalian family and have crania more tightly integrated than their relatives (Hedrick et al 2020). The major axis of cranial variation in Phyllostomidae distinguishes nectivorous and frugivorous species and almost exclusively describes variation in relative facial length, which in turn determines mechanical advantage of the cranium and the most suitable food types (Santana et al 2012, Dumont et al 2014, Hedrick et al 2020, Giacomini et al 2022, Hedrick 2021). Further, facial length dominates evolutionary variation in clade-wise analyses of bats (Hedrick & Dumont 2018, Arbour et al 2019, and see Arbour et al 2021), and heterochrony has been invoked as a primary mechanism generating cranial diversity in bats (Camacho et al 2019, 2020, 2021).

The exploitation of CREA as a line of least resistance seems likely in Chiroptera as in ruminants, and other mammalian clades may similarly deploy simple evolutionary changes in size to achieve adaptation of cranial form through CREA.

## Conclusion

This study builds on a growing body of literature showcasing the profound macroevolutionary implications that ontogenetic allometric patterns may have (e.g., Watanabe 2018, Feiner et al 2021, Navalón et al 2021, Fabbri et al 2021, Chatterji et al 2022, Pavón-Vázquez et al 2022). The findings presented here suggest that morphological diversification of the ruminant skull proceeded along an evolutionary line of least resistance defined by allometry (CREA), and that the eccentricity of population-level variation acted as a facilitator to morphological diversification in that direction because this direction of variation confers adaptive value. This study demonstrates that biases on intraspecific variation at can be reflected in biases on interspecific variation, in this case over 30 million years of ruminant evolution. A key goal of future work should be to understand the ubiquity of exploitation of CREA as a LLR in mammalian evolution, and to contrast the macroevolutionary consequences of CREA in clades where changes in relative facial length have different fitness consequences.

## Acknowledgements

The authors thank Daryl Codron for providing percent-grass-in-diet data, Jessica Maisano for providing the *Saiga tatarica* CT-scan, and Ada Klinkhamer for providing the *Megaloceros giganteus* 3D model. We also greatly appreciate helpful comments on earlier versions of this manuscript by the Jablonski lab at University of Chicago and Angielczyk lab at the Field Museum. We thank Phylopic contributors for the silhouettes of ruminant species used in our figures.

## Funding

This study was supported by the National Science Foundation Graduate Research Fellowship under Grant no. 2020293653.

### Data Availability Statement

All data and R code to replicate these analyses are provided in the supplemental materials and at the lead author’s GitHub page (https://github.com/danielrhoda/ruminant_allometry).

