## Supplemental Materials for "Diversification of the Ruminant Skull Along an Evolutionary Line of Least Resistance"

Landmark definitions:

| | |
|---|---|
| 1 | Suture junction between the two frontals and the nasals; at the frontal end when there is a gap |
| 2 | Suture junction between the two frontals and the parietal; at the frontal end when there is a gap |
| 3 | Supraoccipital boss at the dorsal end of the bone |
| 4 | Anteroventral suture end of the two palate bones |
| 5 | Posteroventral suture end of the two palate bones |
| 6 | Meeting point between the two basioccipitals on the foramen magnum rim |
| 7 | Meeting point between the two occipitals on the foramen magnum rim |
| 8 | Anterior tip of the premaxilla |
| 9 | Infraorbital foramen; the most posterior points on the rim from lateral view |
| 10 | Anterobuccal edge of the third premolar alveoli; meeting point between the alveoli and the tooth |
| 11 | Anterobuccal edge of the first molar alveoli; meeting point between the alveoli and the tooth |
| 12 | Anterodorsal suture end between the jugal and the squamosal; at the point where the suture turns in the medial aspect of the zygomatic arch |
| 13 | Posteroventral suture end between the jugal and squamosal; at the point where the suture turns into the lateral aspect of the zygomatic arch |
| 14 | Suture junction between the parietal, squamosal, and occipital bones |
| 15 | Suture junction between the parietal, squamosal, and alisphenoid bones |
| 16 | Meeting point between the jugal, frontal, and orbital rims on the postorbital bar |
| 17 | Meeting point between the lacrimal, frontal, and orbital rims |
| 18 | Meeting point between the lacrimal, jugal, and orbital rims |
| 19 | Anterior suture end between the lacrimal and the jugal; at the point where the suture turns dorsally |
| 20 | Posterior suture end between the maxilla and the nasal |
| 21 | Subnasal meeting point of the maxilla (sometimes premaxilla) and the nasal |
| 22 | Anterior tip of the nasal |
| 23 | Meeting point between the basioccipital and the foramen magnum condyle |
| 24 | Foramen ovale; the most posterior point on the rim |
| 25 | Posterobuccal edge of the third molar alveoli; meeting point between the alveoli and the tooth |

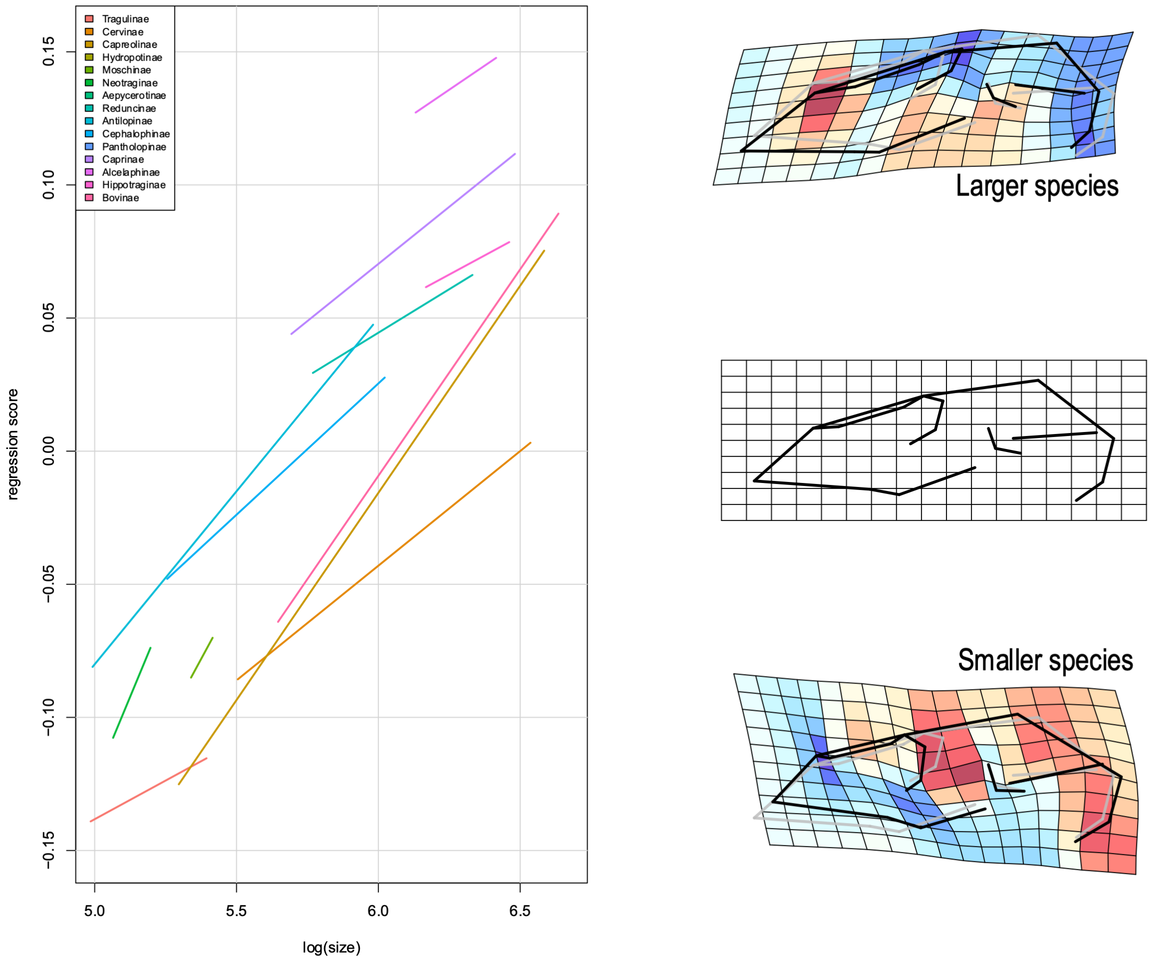


**Supplemental Figure 1**. Homogeneity of slopes test of interspecific allometry in the ruminant skull. Allometric trajectory significantly differed by subfamily, but in each subfamily larger species have proportional longer faces.

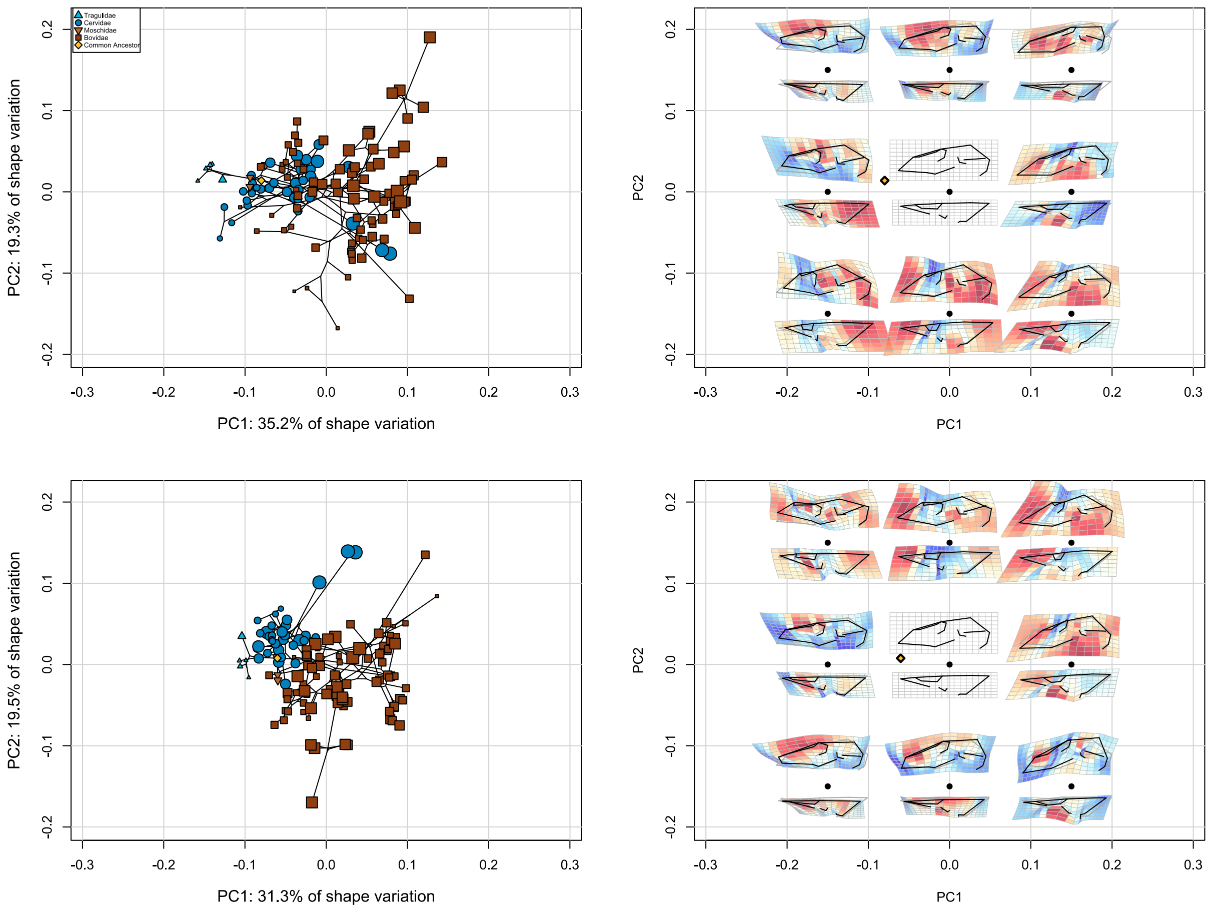


**Supplemental Figure 2**. PCA morphospace with the raw data (top row) and with evolutionary allometry removed (bottom row). Shape variation across the morphospaces are presented in the right column. An interactive dashboard is available to visualize these different ordinations (https://danielrhoda.shinyapps.io/Ruminant_Dashboard/).

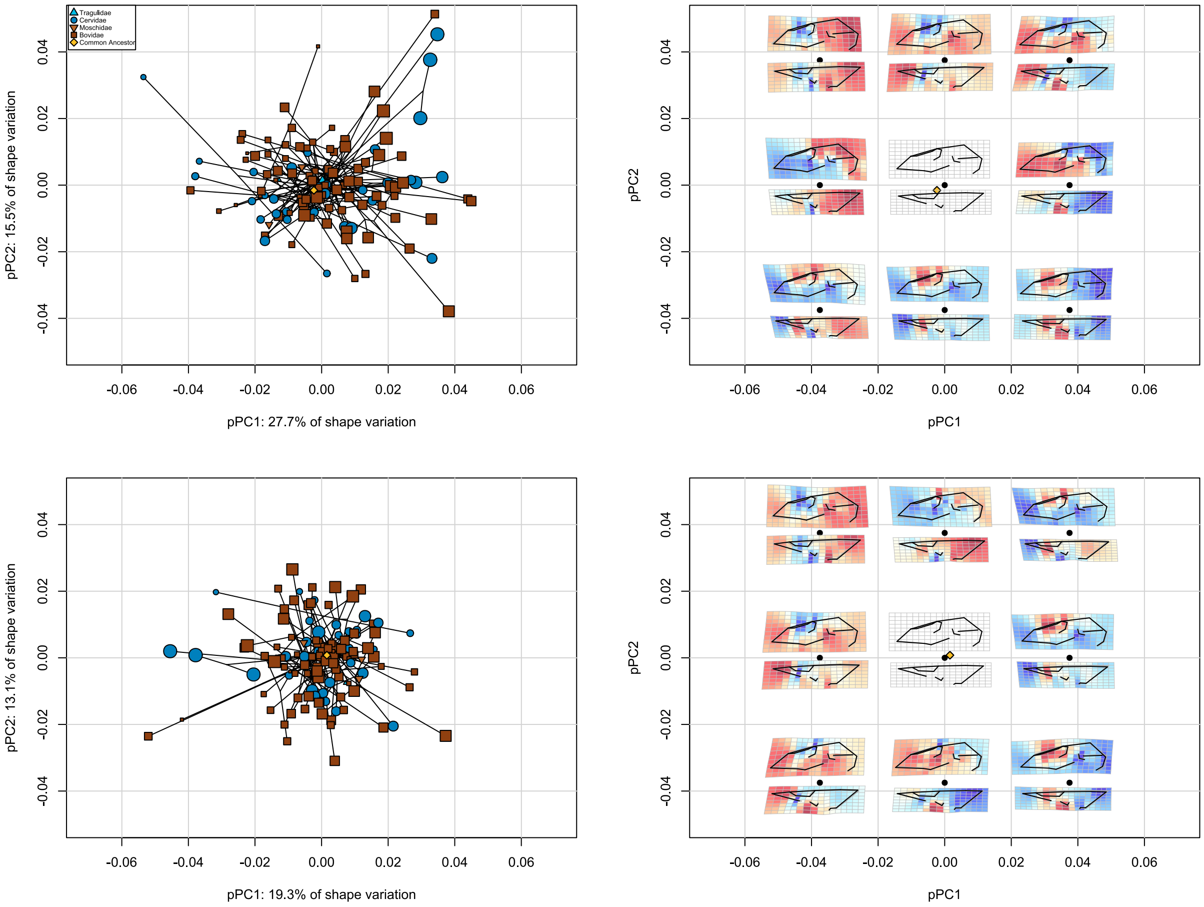


**Supplemental Figure 3**. pPCA morphospace with the raw data (top row) and with evolutionary allometry removed (bottom row). Shape variation across the morphospaces are presented in the right column. An interactive dashboard is available to visualize these different ordinations (https://danielrhoda.shinyapps.io/Ruminant_Dashboard/).

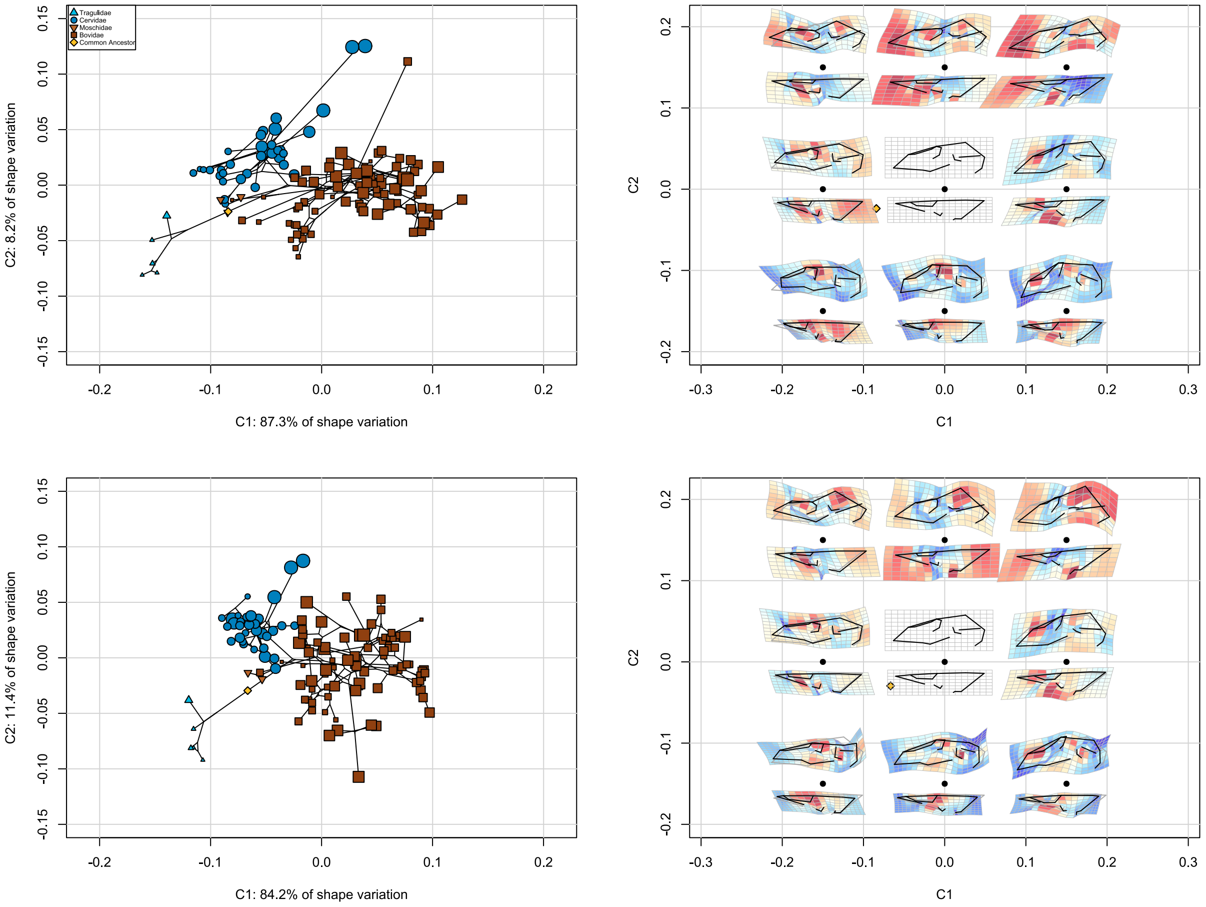


**Supplemental Figure 4**. Phylogenetically-aligned components analysis morphospace with the raw data (top row) and with evolutionary allometry removed (bottom row). Shape variation across the morphospaces are presented in the right column. An interactive dashboard is available to visualize these different ordinations (https://danielrhoda.shinyapps.io/Ruminant_Dashboard/).

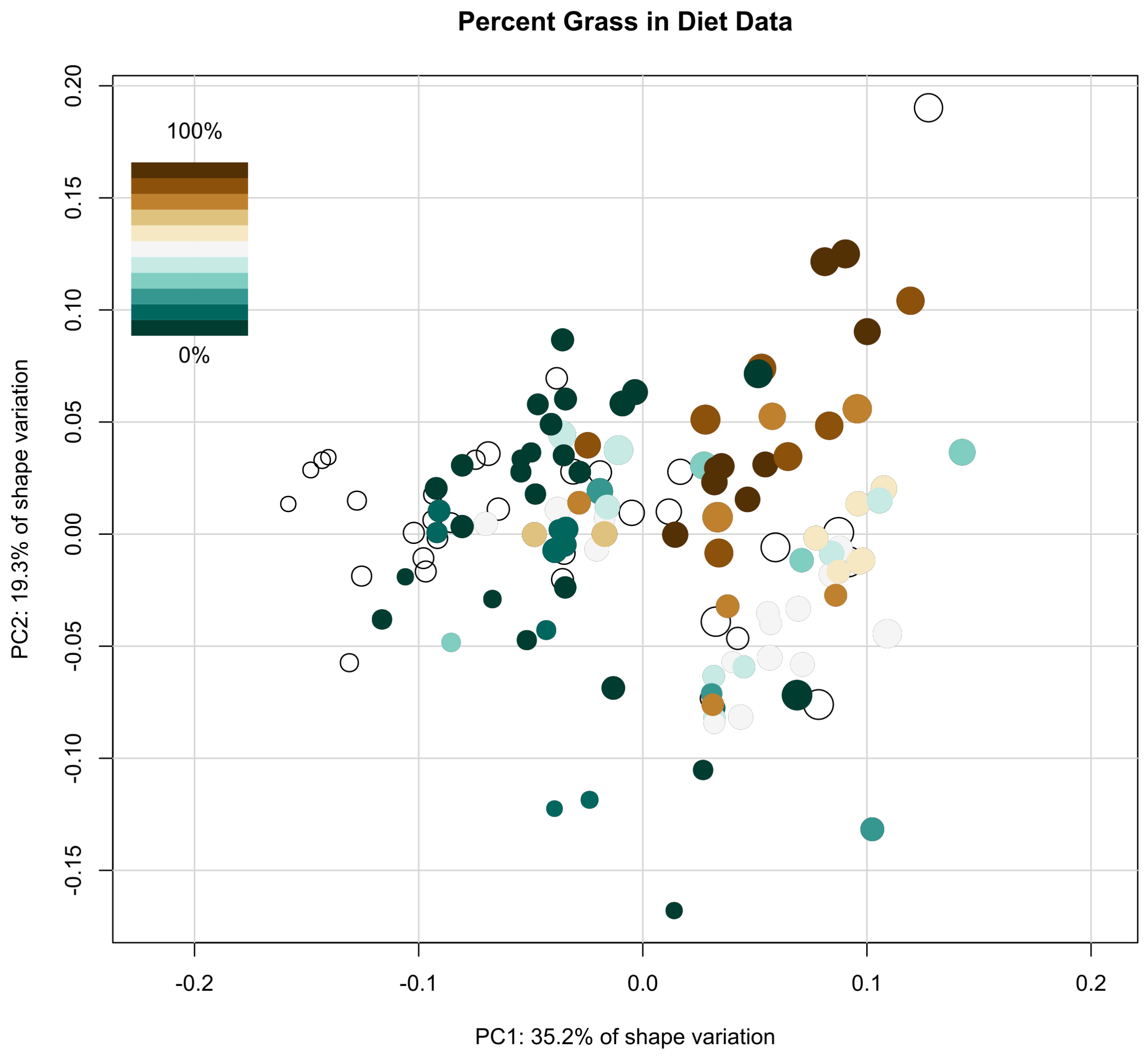


**Supplemental Figure 5**. Interspecific morphospace with species colored by their % grass in diet value, where exclusive grazers have 100% grass in diet. The uncolored points are species for which these data were unavailable.

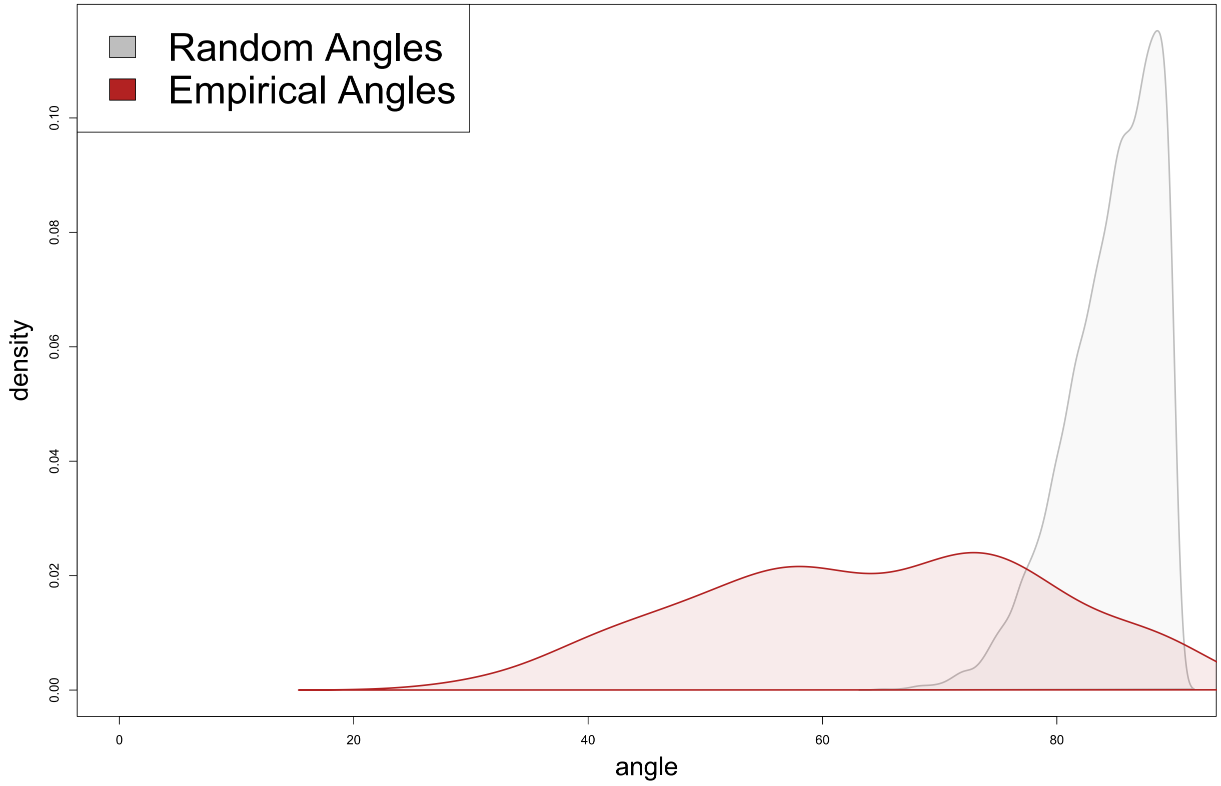


**Supplemental Figure 6.** Distributions of (gray) random angles of the same dimensionality of our dataset (k=75) and (red) observed angles between the direction of divergence and CREA from our empirical dataset. Note that angles here can have a maximum value of 90 degrees.
